# Genetic recombination in disgust-associated bitter taste-responsive neurons of the central nucleus of amygdala in male mice

**DOI:** 10.1101/2020.08.02.233262

**Authors:** Daisuke H. Tanaka, Shusheng Li, Shiori Mukae, Tsutomu Tanabe

**Author notes:** Author for correspondence: Tsutomu Tanabe; Department of Pharmacology and Neurobiology, Graduate School of Medicine, Tokyo Medical and Dental University, 1-5-45 Yushima, Bunkyo-ku, Tokyo 113-8519 Japan.

## Abstract

A bitter substance induces specific orofacial and somatic behavioral reactions such as gapes in mice as well as monkeys and humans. These reactions have been proposed to represent affective disgust, and therefore, understanding the neuronal basis of the reactions would pave the way to understand affective disgust. It is crucial to identify and access the specific neuronal ensembles that are activated by bitter substances, such as quinine, the intake of which induces disgust reactions. However, the method to access the quinine-activated neurons has not been fully established yet. Here, we show evidence that a targeted recombination in active populations (TRAP) method, induces genetic recombination in the quinine-activated neurons in the central nucleus of the amygdala (CeA). CeA is one of the well-known emotional centers of the brain. We found that the intraoral quinine infusion, that resulted in disgust reactions, increased both *cFos*-positive cells and *Arc*-positive cells in the CeA. By using *Arc*-CreER;Ai3 TRAP mice, we induced genetic recombination in the quinine-activated neurons and labelled them with fluorescent protein. We confirmed that the quinine-TRAPed fluorescently-labelled cells preferentially coexpressed *Arc* after quinine infusion. Our results suggest that the TRAP method can be used to access specific functional neurons in the CeA.

## Introduction

The specific orofacial and somatic behavioral reactions, such as gapes and forelimb flails, are observed in many mammals including humans, monkeys, and rodents in response to intraoral infusion of a bitter substance [1,2]. These reactions have been proposed to reflect affective disgust, rather than a sensory reflex, consummatory behaviors, or an avoidance motivation [2,3]. Thus, an understanding of the neural basis of these behavioral reactions (disgust reactions) would provide important insights for understanding affective disgust.

The disgust reactions are associated with cFos expression, a marker of neuronal activity, in several brain regions such as the external part of the medial parabrachial nucleus, nucleus of the solitary tract [4], rostral part of the central nucleus of the amygdala (CeA), insular cortex [5] and interstitial nucleus of the posterior limb of the anterior commissure (IPAC) [6]. These brain regions appear to consist of functionally heterogeneous neurons [7,8,9–11]. Therefore, the cFos-positive neurons associated with the disgust reactions are likely to constitute only a fraction of neurons in each brain region. Indeed, the *cFos*-positive neurons and the neurons expressing *Arc*, another marker of neuronal activity [12,13], that are associated with the disgust reactions, constitute a subset of neurons in the IPAC [6]. To investigate the physiological role of the neurons whose activity is associated with the disgust reactions, it would be crucial to establish a method to specifically access the disgust-associated neurons. While the quinine-activated disgust-associated neurons in the IPAC are preferentially accessible by a targeted recombination in active populations (TRAP) method [6,14], it remains unclear whether the accessibility is restricted to the IPAC or not. It is interesting to investigate whether the quinine-activated neurons in the CeA [5] are accessible by TRAP because of the following reasons: (1) the CeA is one of the well-known emotional centers but its role in the disgust reactions remains unclear; (2) the CeA consists of functionally heterogeneous neurons, and the expression of a single molecular marker and Cre transgenics in the molecularly defined neurons likely cannot capture the functional heterogeneity in the CeA [10,15,16]; (3) the CeA neurons are not successfully TRAPed by any stimuli [14,17,18,19]; (4) the CeA and the IPAC, where a significant number of neurons were successfully TRAPed by quinine infusion [6], are directly continuous anatomically and are grouped as a part of the distinct anatomical division, the central division of the extended amygdala [7].

In the present study, we set out to genetically access and label the quinine-activated disgust-associated neurons in the CeA by the TRAP method.

## Material and methods

### Subjects

The histological slices analyzed for the current study were from the same brain tissues from an earlier report examining the IPAC [6]. In brief, wild-type C57BL/6J mice were obtained from Japan SLC, Inc. *Arc-CreER* mice (JAX stock #021881) [14] or *cFos-CreER* mice (stock #021882) [14] were crossed with *Ai3*(*RCL-EYFP*) (*Ai3*) mice (stock #007903) [20] to produce *Arc-CreER;Ai3* mice or *cFos-CreER;Ai3* mice, respectively. All the animal experiments were approved (No. 0150384A, 0160057C2, 0170163C, A2017-194A, A2018-138C4) by the Institutional Animal Care and Use Committee of Tokyo Medical and Dental University and performed in accordance with the relevant guidelines and regulations.

### Surgery

The surgery for implantation of intraoral tube was performed as described previously [6]. Briefly, a mixture of midazolam (4 mg/kg body weight (BW)), butorphanol (5 mg/kg BW), and medetomidine (0.3 mg/kg BW) was used to anesthetize mice. A curved needle attached to an intraoral polyethylene tube was inserted from the incision site and advanced subcutaneously posterior to the eye to exit at a point lateral to the first maxillary molar on the right side of the mouth.

### Stimulation for fluorescent in situ hybridization (FISH) experiments

The stimulation for FISH experiments (Fig. 2) was performed as described previously [6]. In brief, mice received intraoral infusions of water, 5.4 mM saccharin solution, or 3 mM quinine solution. Each infusion was 50 μL in volume and lasted for 1 min, and the two infusions were performed with a 1 min interval. Five minutes after completion of the last infusion, mice were deeply anesthetized and perfused with 4% paraformaldehyde in phosphate-buffered saline.

### Dual stimulation for TRAP and FISH

The dual stimulation for TRAP and FISH (Fig. 3, 4) was performed as described previously [6]. In brief, mice received intraoral infusions of 5 μL of water (Water-TRAP) or 30 mM quinine solution (Quinine-TRAP) 12 times with 5 min intervals. At 1-1.5 h after the last infusion, mice were injected intraperitoneally (i.p.) with 20 mL/kg 4-hydroxytamoxifen (4-HT) solution. Five to seven days after the 4-HT injection, all mice received intraoral infusions of 5 μL of 30 mM quinine solution twice, with 1 min interval, and then deeply anesthetized and perfused five minutes after completion of the last infusion (Quinine-fix). The reason of using these stimulation parameters for TRAP was described previously [6].

### Taste reactivity test during stimulation for TRAP

The taste reactivity test during intraoral water or quinine infusions for TRAP and scoring were performed as described previously [6] with minor modifications. Briefly, the orofacial and somatic behavior of mice was video recorded for 1 min after 1st and 7th water or quinine infusions (2 min in total). Liking and disgust taste reactivity patterns during intraoral solution infusion were scored manually frame-by-frame (30 frames/sec) [2,21] by a blind observer [6].

### Immunofluorescence staining

The immunofluorescence staining was performed as described previously [6]. Briefly, the sections were incubated 1-3 overnights at 4°C in the following primary antibodies: goat anti-choline acetyltransferase (1:200; Millipore), rabbit anti-substance P (1:500; Immunostar), chicken anti-GFP (1:500; abcam), mouse anti-enkephalin (1:400; Millipore), and/or biotin-conjugated mouse anti-NeuN (1:500; Chemicon). After washing, the sections were incubated in the secondary antibodies and then incubated with Hoechst33258.

### FISH in combination with Immunofluorescence

FISH in combination with Immunofluorescnece was performed as described previously [6]. In brief, the cryosections on slides were incubated with digoxigenin-labeled *cFos* or *Arc* riboprobes [22] overnight at 65 °C. Following washes, sections were incubated with the primary antibodies described above for 3 h. After washes, sections were incubated with alkaline phosphatase-conjugated sheep anti-digoxigenin antibody (1:2000; Roche) and the secondary antibodies overnight at 4°C. After washes, sections were incubated with HNPP/Fast Red TR solution (Roche) for 20-30 min and then incubated with Hoechst33258.

### Cell quantification

Since a rostrocaudal gradient characterized the quinine-stimulated cFos response, with the greatest number of labeled cells situated rostrally in the CeA [5], the rostral and middle levels of the CeA along the rostrocaudal axis, which corresponds to the CeA from Bregma level −0.70 to −1.46 mm [23], were analyzed. Sections were collected from Bregma level −0.70 to −1.46 mm with 0.1-0.3 mm intervals and all sections without tissue damage and/or distortion around the CeA on slides were stained and analyzed.

NeuN-, *cFos*-, *Arc*-, and YFP-positive cells were counted manually by a blind observer, and Hoechst33258-positive cells were counted with NIH ImageJ software (version 1.40 g).

To calculate the ratio of the number of NeuN-positive cells to the number of Hoechst33258-positive cells (Fig. 1D), the number of NeuN-positive cells was divided by the number of Hoechst33258-positive cells (Table 1).

**Fig. 1.**
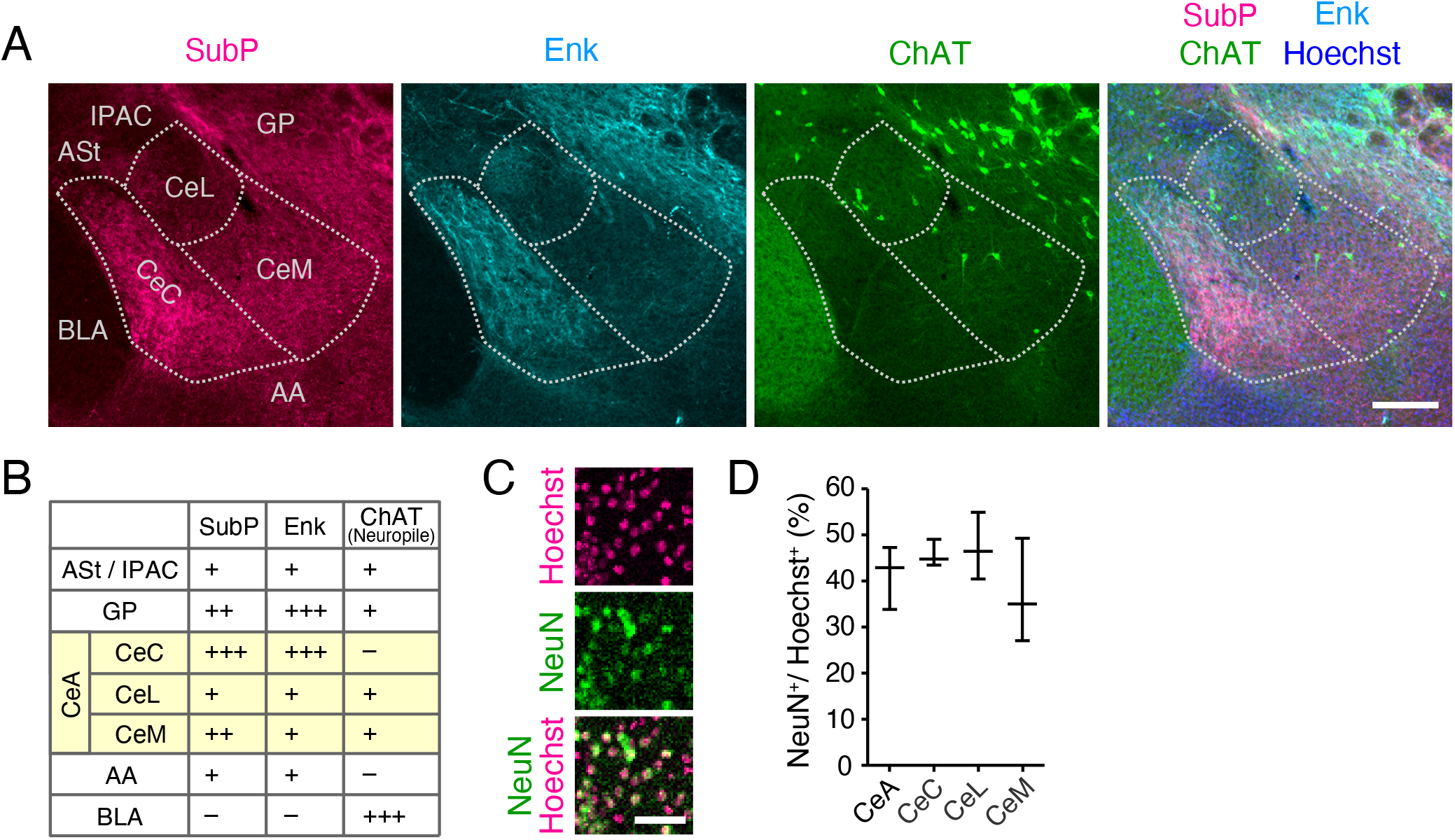
The expressions of substance P (SubP), enkephalin (Enk) and choline acetyltransferase (ChAT) distinguish the subdivisions of the central nucleus of the amygdala (CeA) from adjacent brain regions. (**A**) Representative images of SubP (magenta), Enk (light blue), ChAT (green) expressions and Hoechst33258 staining (blue) in the central nucleus of the amygdala (white dotted lines) and the adjacent regions of wild-type mice. (**B**) Summary of the signal intensity of SubP, Enk and ChAT (neuropile) in the CeA and adjacent regions. Note that, as for ChAT, neuropile-like signals, but not cell body-like signals, were focused. Plus signs, +++, ++ and +, mean that the relative signal intensities were high, middle and low, respectively. Minus sign, -, means signals undetectable. (**C**) Representative images of NeuN expression (green) and Hoechst33258 staining (magenta) in the CeA. (**D**) The proportion of the number of NeuN-positive cells to the number of all cells (Hoechst33258-positive cells) in the subregions of the CeA (*n* = 5 sections, 3 brains). ASt, Amygdalostriatal transition area; IPAC, interstitial nucleus of the posterior limb of the anterior commissure; GP, globus pallidus; CeC, central amygdaloid nucleus capsular part; CeL, central amygdaloid nucleus lateral division; CeM, central amygdaloid nucleus medial division; AA, anterior amygdaloid area; BLA, basolateral amygdaloid nucleus anterior part. Scale bars: 200 *μ*m in **A**; 50 *μ*m in **C**.

**Table 1.** Cell numbers counted in quantitative analyses.

| Fig.1D | NeuN+ | Hoechst+ |
| --- | --- | --- |
| CeA | 3202 | 7544 |
| CeC | 1091 | 2376 |
| CeL | 623 | 1273 |
| CeM | 1488 | 3895 |

| Fig.2C,D |  | *cFos*+ | Hoechst+ | (NeuN+) | *Arc*+ | Hoechst+ | (NeuN+) |
| --- | --- | --- | --- | --- | --- | --- | --- |
| CeA | Water | 52 | 4628 | 1913 | 81 | 14015 | 5794 |
|  | Saccharin | 109 | 9824 | 4062 | 88 | 16164 | 6683 |
|  | Quinine | 244 | 7126 | 2946 | 393 | 10265 | 4244 |
| CeC | Water | 28 | 1755 | 750 | 52 | 4751 | 2032 |
|  | Saccharin | 53 | 3868 | 1654 | 46 | 5342 | 2284 |
|  | Quinine | 125 | 2910 | 1244 | 193 | 3553 | 1519 |
| CeL | Water | 4 | 881 | 375 | 17 | 2304 | 981 |
|  | Saccharin | 20 | 1751 | 746 | 21 | 2689 | 1145 |
|  | Quinine | 48 | 1283 | 546 | 168 | 2289 | 975 |
| CeM | Water | 20 | 1992 | 740 | 12 | 6960 | 2584 |
|  | Saccharin | 36 | 4205 | 1561 | 21 | 8133 | 3019 |
|  | Quinine | 71 | 2933 | 1089 | 32 | 4423 | 1642 |

| Fig.4A,B |  | YFP+ | Arc+ | Hoechst+ | (NeuN+) |
| --- | --- | --- | --- | --- | --- |
| CeA | Water-TRAP | 146 | 361 | 19729 | 8157 |
|  | Quinine-TRAP | 258 | 161 | 16683 | 6897 |
| CeC | Water-TRAP | 57 | 222 | 6634 | 2837 |
|  | Quinine-TRAP | 123 | 100 | 5793 | 2477 |
| CeL | Water-TRAP | 60 | 112 | 4539 | 1933 |
|  | Quinine-TRAP | 103 | 55 | 3786 | 1612 |
| CeM | Water-TRAP | 29 | 27 | 8556 | 3177 |
|  | Quinine-TRAP | 32 | 6 | 7104 | 2637 |

To calculate the ratio of the number of *cFos*- (Fig. 2C, Table 1), *Arc*- (Fig. 2D, 4, Table 1), or YFP-positive cells (Fig. 4, Table 1) to the number of NeuN-positive cells, the number of NeuN-positive cells in the subregion was firstly calculated by multiplying the number of Hoechst33258-positive cells by the pre-determined ratio of the number of NeuN-positive cells to that of Hoechst33258-positive cells (Fig. 1D). Then, the number of *cFos*-, *Arc*-, or YFP-positive cells was divided by the calculated number of NeuN-positive cells.

**Fig. 2.**
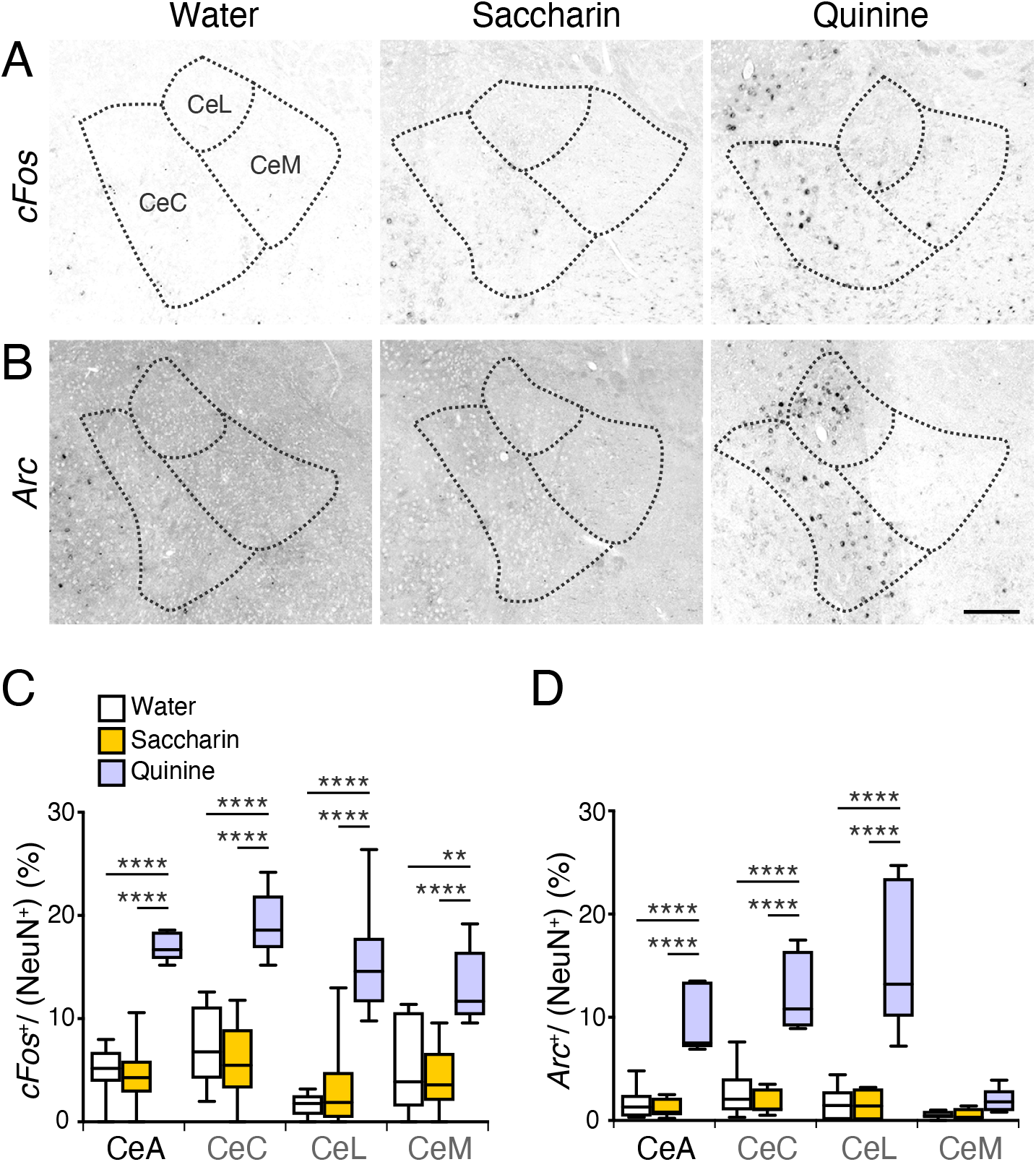
Quinine infusion induces both *cFos* and *Arc* expression in the CeA. (**A, B**) Representative images of *cFos* (**A**) and *Arc* (**B**) expression (black) around the CeA (black dotted line) in wild-type mice after intraoral infusion of water (left panels), saccharin solution (middle panels) or quinine solution (right panels). (**C**) The ratio of the number of *cFos*-positive cells to the number of NeuN-positive cells in the subregions of the CeA from mice after intraoral infusion of water (*n* = 10 sections, 6 mice from 3 independent experiments), saccharin solution (*n* = 15 sections, 8 mice from 4 independent experiments) or quinine solution (*n* = 12 sections, 6 mice from 4 independent experiments). (**D**) The ratio of the number of *Arc*-positive cells to the number of NeuN-positive cells in the subregions of the CeA from mice after intraoral infusion of water *(n* = 10 sections, 6 mice from 3 independent experiments), saccharin solution (*n* = 12 sections, 7 mice from 3 independent experiments) or quinine solution (*n* = 7 sections, 5 mice from 3 independent experiments). (NeuN+) represents the calculated number of NeuN-positive cells. \*\**P* < .01, \*\*\*\**P* < .0001 (Tukey’s test).

To calculate TRAP specificity (Fig. 4C), the number of YFP;*Arc* double-positive cells was divided by the number of YFP-positive cells.

To calculate TRAP efficiency (Fig. 4D), the number of YFP;*Arc* double-positive cells was divided by the number of *Arc*-positive cells.

To calculate the chance ratio of the number of YFP;*Arc* double-positive cells to the number of NeuN-positive cells (Fig. 4F,G), the number of NeuN-positive cells was firstly calculated as mentioned above, and the ratio of the number of YFP-positive cells to the number of NeuN-positive cells was multiplied by the ratio of the number of *Arc*-positive cells to the number of NeuN-positive cells.

To calculate TRAP preference (Fig. 4E), the real ratio of the number of YFP;*Arc* double-positive cells to the number of NeuN-positive cells was divided by the chance ratio of the number of YFP;*Arc* double-positive cells to the number of NeuN-positive cells. Thus, the value 1 in TRAP preference means that the overlap between YFP and *Arc* can be explained by chance.

### Statistical analysis

All statistics were performed using GraphPad Prism 8 software (GraphPad Software). Independent sampling (i.e. one data point per animal) was used for all statistical analysis. Differences in multiple parameters between the three groups were analyzed by two-way ANOVA followed by Tukey’s multiple comparison test (Fig. 2C,D). Differences in multiple parameters between the two groups were analyzed by two-way ANOVA followed by Sidak’s multiple comparison test (Fig. 3B,4). A p value < 0.05 was considered statistically significant. All tests were two-tailed. Data were expressed as boxes representing 25-75^th^ percentiles, whiskers representing minimum-max, and lines representing medians.

**Fig. 3.**
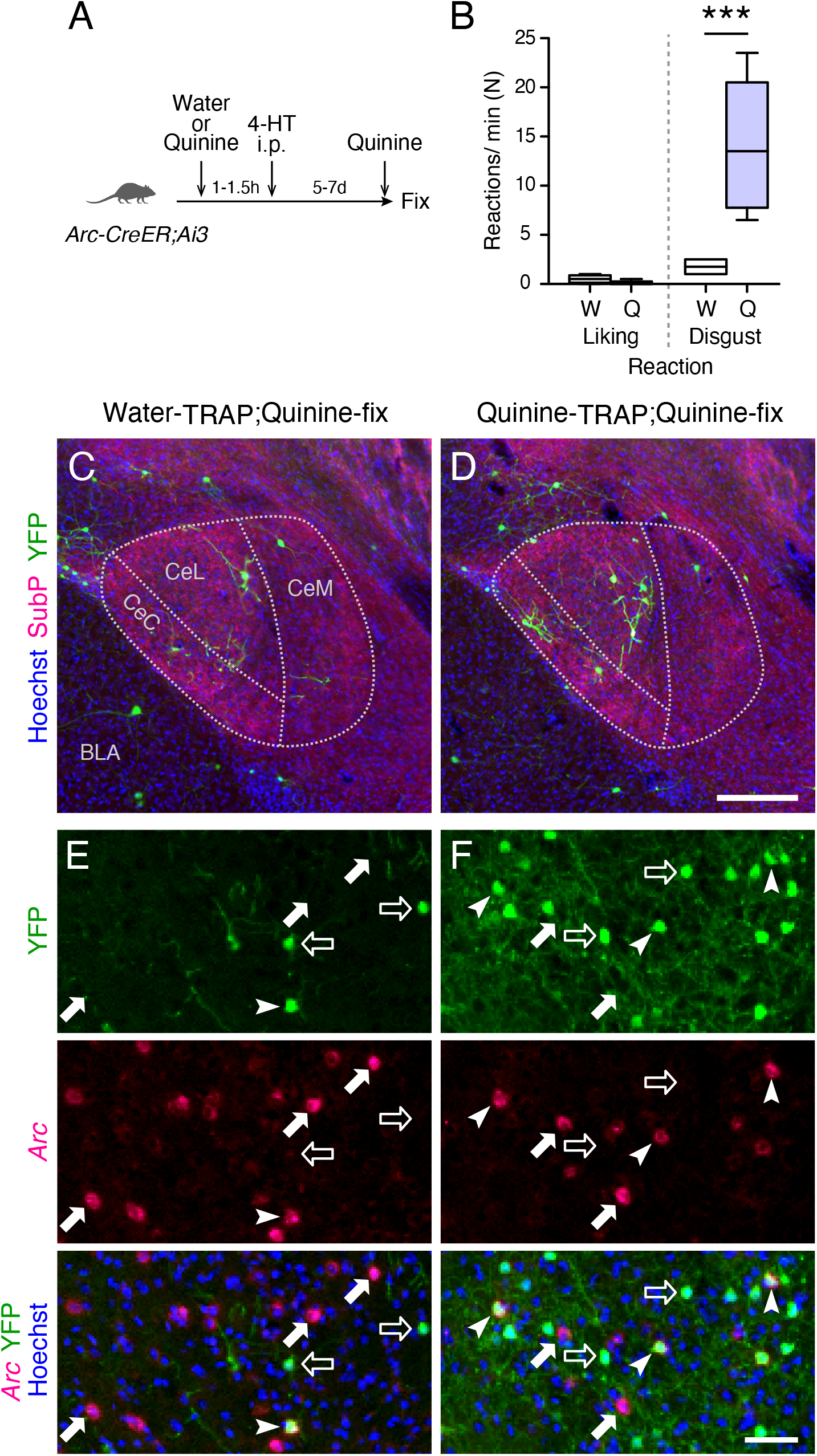
Quinine-TRAP in the CeA of *Arc-CreER;Ai3* mice. (**A**) Schematic of experimental design. (**B**) The behavioral liking and disgust reaction of *Arc-CreERT;Ai3* mice in response to intraoral water (*n* = 4 mice) or quinine (*n* = 5 mice) infusion. \*\*\**P* < .001 (Sidak’s test). (**C, D**) Representative images of YFP-positive cells (green) around the CeA (white dotted lines) of *Arc-CreER;Ai3* mice after Water-TRAP (**C**) or Quinine-TRAP (**D**). (**E**, **F**) Representative images of overlap of YFP with *Arc* in the CeA of *Arc-CreER;Ai3* mice after Water-TRAP (**E**) or Quinine-TRAP (**F**). YFP single-positive cells (open arrows), *Arc* single-positive cells (filled arrows) and YFP;*Arc* double-positive cells (arrowheads) were observed. i.p., intraperitoneal; 4-HT, 4-hydroxytamoxifen; W, water; Q, quinine. Scale bars: 200 *μ*m in **C**, **D**; 50 *μ*m in **E**, **F**.

## Results

### The subdivisions of the CeA could be distinguished from adjacent brain regions based on differential expression of several molecules

We first examined molecular markers to distinguish three subdivisions of the CeA, namely the central amygdaloid nucleus capsular part (CeC), lateral division (CeL), and medial division (CeM) [15] from adjacent brain regions. Consistent with previous reports [24,25], substance P, enkephalin and choline acetyltransferase were differentially expressed around the CeA and its subdivisions. For example, the signal intensities of substance P and enkephalin were higher in the CeC, compared to adjacent regions such as the CeL, CeM and basolateral amygdala (Fig. 1A,B). The intensity of choline acetyltransferase signal was higher in the basolateral amygdala than the adjacent CeA (Fig. 1A,B). Thus, we used these markers in the following anatomical experiments to determine the CeA region and its subdivisions.

We also counted the number of both NeuN-positive cells (neurons) and Hoechst-positive cells (all types of cells including neurons) in each subdivision of the CeA. We then calculated the ratio of the number of NeuN-positive cells to that of Hoechst-positive cells. We found that ~40% of Hoechst-positive cells were positive for NeuN in all three subdivisions of the CeA (Fig. 1C,D). This ratio was used in the following experiments to calculate the number of NeuN-positive cells based on the number of Hoechst-positive cells by multiplying the ratio by the number of Hoechst-positive cells (for details see Material and methods).

### Quinine infusion associated with disgust reactions increased both *cFos*-positive cells and *Arc*-positive cells in the CeC and CeL

In the previous study, we examined the taste reactivity in response to intraoral infusion of water, saccharin, or quinine in wild-type mice [6]. Quinine infusion significantly induced disgust reactions compared to water or saccharin infusion [6]. The brain cryosections of the mice, fixed five minutes after the taste reactivity test [6], were stained for *cFos* (Fig. 2A) and *Arc* (Fig. 2B) mRNA by fluorescent *in situ* hybridization (FISH). The proportion of *cFos*-positive cells (*cFos* cells) and *Arc*-positive cells (*Arc* cells) to NeuN-positive cells (neurons) were significantly higher in the CeA of the quinine-infused mice, compared to water-infused or saccharin-infused mice (Fig. 2C,D), although only few *Arc* cells were observed in the CeM (Fig. 2B,D). Thus, quinine infusion associated with disgust reactions significantly increased both *cFos* cells and *Arc* cells in the CeC and CeL.

### Quinine-activated disgust-associated neurons were preferentially TRAPed in the CeC and CeL of *Arc-CreER;Ai3* mice

To test whether the disgust-associated neurons in the CeC and CeL are accessible by TRAP, we used *Arc-CreER;Ai3* double-transgenic mice, in which neuronal activation would induce cytoplasmic expression of CreER^T2^ that enters the nucleus by binding with 4-HT and causes recombination, resulting in permanent YFP expression (TRAPed) in the activated neurons [6]. Quinine or water was infused 1-1.5 h before 4-HT intraperitoneal injection, while quinine was infused in all of the mice 5 min before fixation (Fig. 3A) [6]. We first confirmed that the *Arc-CreER;Ai3* mice showed significant disgust reactions in response to intraoral quinine infusion (Fig. 3B). TRAP following quinine infusion (Quinine-TRAP) induced significantly more YFP-positive cells (YFP cells or TRAPed cells) in the CeC and CeL (Fig. 3D, 4A), compared to a water-infused control (Water-TRAP) (Fig. 3C, 4A). Quinine-TRAP resulted in few YFP cells in the CeM (Fig. 4A) which is in consistent with the fact that few *Arc* cells were found in the CeM after quinine infusion (Fig. 2B,D).

To test whether Quinine-TRAP in the CeC and CeL preferentially labels quinine-activated neurons, we examined the overlap between YFP and *Arc* mRNA (Fig. 3E,F). TRAP specificity (YFP;*Arc* double-positive cells among YFP cells) was not significantly different between Quinine-TRAP and Water-TRAP (Fig. 4C), potentially due to the tendency of less *Arc* cells in Quinine-TRAP compared to Water-TRAP (Fig. 4B). On the other hand, TRAP efficiency (YFP;*Arc* double-positive cells among *Arc* cells) was significantly higher in Quinine-TRAP compared to Water-TRAP (Fig. 4D). The overlap ratio between YFP and *Arc* was significantly higher in Quinine-TRAP than by chance [YFP^+^/NeuN^+^ (Fig. 4A) × *Arc*^+^/NeuN^+^ (Fig. 4B)] (Fig. 4G), while that was not found to be significantly different from chance in Water-TRAP (Fig. 4F). TRAP preference [the real ratio of the number of YFP;*Arc* double-positive cells to the number of NeuN-positive cells (Fig. 4F, open boxes; Fig. 4G, purple boxes) was divided by the chance ratio of the number of YFP;*Arc* double-positive cells to the number of NeuN-positive cells (Fig. 4F, 4G, gray boxes)] was significantly higher in Quinine-TRAP compared to Water-TRAP in the CeA as a whole (Fig. 4E). These data suggest that Quinine-TRAP in *Arc-CreER;Ai3* mice preferentially accesses and labels quinine-activated neurons in the CeC and CeL.

**Fig. 4.**
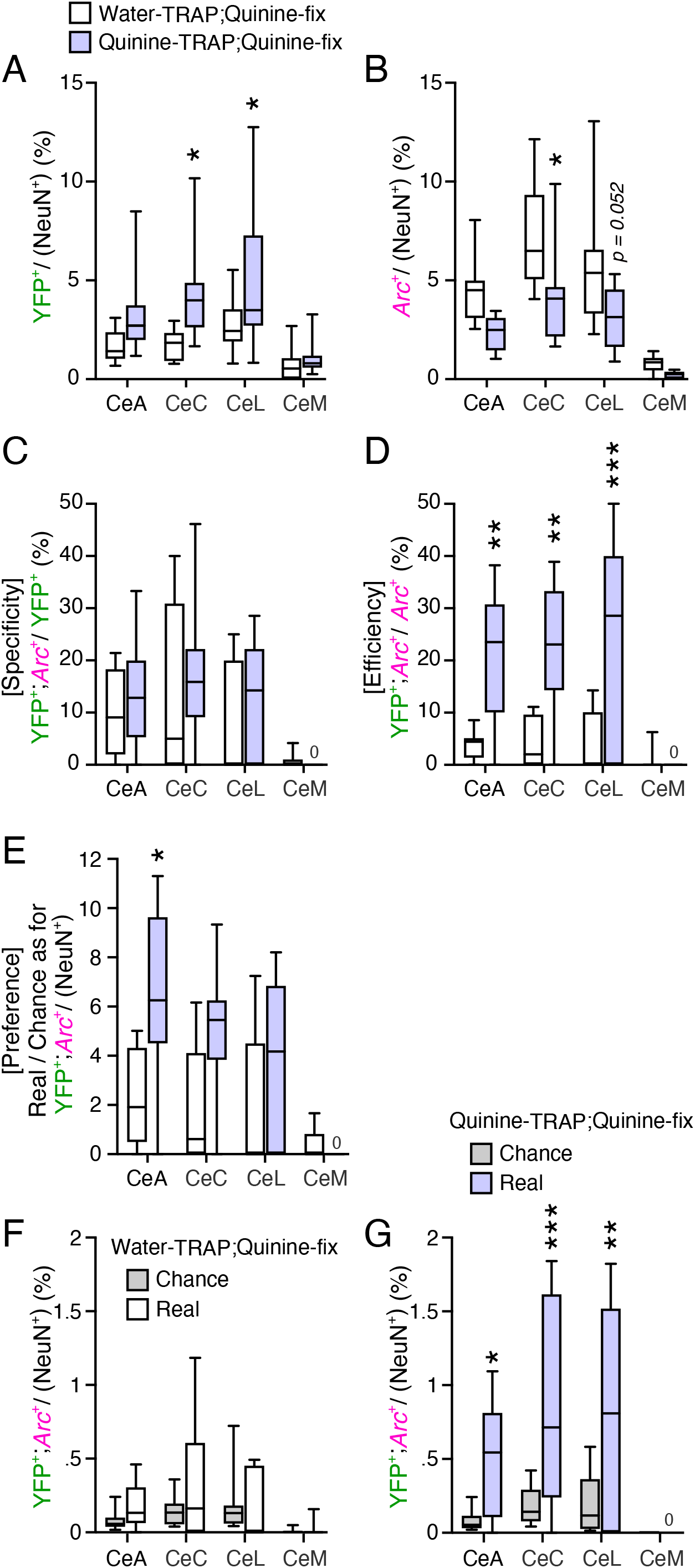
Quinine-TRAP preferentially labels quinine-activated neurons in the CeA. Water-TRAP;Quinine-fix mice (open boxes) (*n* = 33 sections, 9 mice from 4 independent experiments) or Quinine-TRAP;Quinine-fix mice (light purple boxes) (*n* = 23 sections, 7 mice from 3 independent experiments) were analyzed. (**A**) The proportion of the number of YFP-positive cells to the number of NeuN-positive cells. (**B**) The proportion of the number of *Arc*-positive cells to the number of NeuN-positive cells. (**C**) The proportion of the number of YFP;*Arc* double-positive cells to the number of YFP-positive cells, indicating TRAP specificity. (**D**) The proportion of the number of YFP;*Arc* double-positive cells to the number of *Arc*-positive cells, indicating TRAP efficiency. (**E**) The ratio of the real ratio of the number of YFP;*Arc* double-positive cells to the number of NeuN-positive cells to the chance ratio of the number of YFP;*Arc* double-positive cells to the number of NeuN-positive cells (**F, G**), indicating TRAP preference. (**F, G**) The proportion of the number of *YFP;Arc* double-positive cells to the number of NeuN-positive cells for Water-TRAP;Quinine-fix (**F**) or Quinine-TRAP;Quinine-fix (**G**). \**P* < .05, \*\**P* < .01, \*\*\**P* < .001 (Sidak’s test).

We also used *cFos-CreER;Ai3* double-transgenic mice [6], but Quinine-TRAP in this mice failed to induce significantly more YFP cells in the CeA [1.7 ± 0.4% (mean ± s.e.m.) among all neurons, *n* = 8 mice], compared to Water-TRAP (0.9 ± 0.2%, *n* = 6 mice, *P* = 0.804, Sidak’s test).

## Discussion

### Differential expression of *cFos* and *Arc* in the CeM

*cFos* and *Arc* have been well characterized and widely used as markers of neuronal activity in histological sections for decades [12,13,26]. Accumulating evidence suggests that these genes can be differentially regulated in different cell types in distinct brain areas. In the striatum, *Arc* rather than *cFos* appears to be strongly expressed by GABAergic medium spiny neurons [14,27]. In contrast, *cFos* rather than *Arc* appears to be strongly expressed in the thalamic areas [13,28] and in the excitatory granule cells in the cerebellum [14]. We found that intraoral quinine infusion significantly increased *cFos* cells but not *Arc* cells in the CeM (Fig. 2C,D), suggesting that these two activity-dependent genes were differentially regulated in a subset of CeM neurons. There are two possible explanations for the differential expression between *cFos* and *Arc* in the CeM: (1) CeM neurons were activated by quinine; *cFos* expression was induced since neurons were activated; *Arc* expression was not induced by unknown reason even though neurons were activated; (2) CeM neurons were not activated by quinine; *cFos* expression was induced through activity-independent mechanisms [29,30]; *Arc* expression was not induced since neurons were not activated. More direct monitoring of CeM neuronal activity by using electrophysiological recording or Ca^2+^ imaging would be helpful to distinguish these possibilities.

### Genetic access to quinine-activated neurons in the CeA

The CeA orchestrates a diverse set of adaptive behaviors and consists of functionally as well as genetically defined cells types that control specific behavioral outputs. Defining cellular identity based on just the expression of a single molecular marker, likely cannot capture the functional heterogeneity within molecularly identified populations [10]. For example, while PKC-delta is expressed in a~70% of cFos cells in the CeL in response to intraoral quinine infusion, the PKC-delta and cFos double-positive cells constitute only ~10% of whole PKC-delta-positive cells in the CeL [31]. Under water-deprived condition, spontaneous quinine intake induces *cFos* in only ~5% and ~30% of PKC-delta-positive cells in the CeL and CeC, respectively [15]. Thus, PKC-delta could not be used as a perfect marker of quinine-stimulated cFos cells in the CeL and CeC, which prompted us to test the TRAP method to access the quinine-activated neurons.

Quinine-TRAP successfully labelled more cells compared to Water-TRAP in the CeC and CeL (Fig. 3C,D, 4A). Quinine-TRAP efficiency was ~25% in these regions (Fig. 4D), suggesting that ~25% of quinine-activated cells were TRAPed. This ratio is largely comparable with those of some previous TRAP studies: foot shock stimulation TRAPed ~20-40% of foot shock-activated cells in medial prefrontal cortex, the BLA and the nucleus accumbens [17]; pheromone stimulation TRAPed ~15% of the pheromone-activated cells in the ventromedial hypothalamus [18]. In addition, we confirmed that the Quinine-TRAPed cells were preferentially activated by quinine stimulation (Fig. 4E,G). However, TRAP specificity was not found to be significantly different between Quinine-TRAP and Water-TRAP in the CeA, while it is significantly different in the IPAC [6]. One possible explanation for the low Quinine-TRAP specificity, is that the number of *Arc* cells tended to be less in the Quinine-TRAP compared to Water-TRAP (Fig. 4B). This reduction of *Arc* cells in the Quinine-TRAP has not been observed in the IPAC [6]. Repeated quinine stimulations during TRAP might reduce *Arc* expression in the CeA in the last stimulation just before fixation (compare open boxes and purple boxes in Fig. 4B), resulting in less YFP and *Arc* double-positive cells and thus the lower calculated value of specificity (Fig. 4C). It remains unclear whether the smaller number of *Arc* cells in Quinine-TRAP of the CeA (Fig. 4B) reflects reduced *Arc* expression efficiency in a certain neuronal activity, reduced neuronal activity, or both. However, it would be interesting to identify the cause of the *Arc* differential expression between CeA and IPAC [6].

Although Quinine-TRAP preferentially labelled quinine-activated neurons, it remains unclear whether the Quinine-TRAPed cells encode bitterness of quinine, behavioral disgust reactions, or disgust-related psychological functions. In addition, we cannot exclude the possibility that Quinine-TRAPed cells encode not only disgust-related responses but also other psychological functions such as mental helplessness after inescapable repeated stress by high concentration of quinine. Since TRAP is based on recombination, it would be possible to specifically express genes of interest in the Quinine-TRAPed cells. By combining with Cre-dependent optogenetic/chemogenetic tools, it would be possible to specifically manipulate the activity of TRAPed cells [18] in the CeA to understand their physiological roles in disgust reactions and conscious disgust [32].

## Conclusions

We conclude that quinine-activated disgust-associated neurons in the CeC and CeL are preferentially recombined and genetically labelled by TRAP.

## Abbreviations

ASt: Amygdalostriatal transition area;
IPAC: interstitial nucleus of the posterior limb of the anterior commissure;
GP: globus pallidus;
CeC: central amygdaloid nucleus capsular part;
CeL: central amygdaloid nucleus lateral division;
CeM: entral amygdaloid nucleus medial division;
AA: anterior amygdaloid area;
BLA: basolateral amygdaloid nucleus anterior part;
TRAP: Targeted Recombination in Active Populations;
FISH: fluorescent in situ hybridization;
GFP: green fluorescent protein;
YFP: yellow fluorescent protein;
BW: body weight;
4-HT: 4-hydroxytamoxifen;
i.p.: intraperitoneal;
SubP: substance P;
Enk: enkephalin;
ChAT: choline acetyltransferase;
Hoechst: Hoechst33258;
W: water;
Q: quinine

## Acknowledgements

We thank Drs. M. Kobayashi and N. Mizoguchi at Nihon University for technical advice for intraoral tubing surgery, Dr. H. Takebayashi at Niigata University for *cFos* and *Arc* plasmids. This work was supported by JSPS KAKENHI [grant numbers JP26890011, JP16K07024 and JP19K06938], Takeda Science Foundation and TOBE MAKI Scholarship Foundation to D.H.T.

